# Variability in HIV-1 Transmitted/Founder Virus Susceptibility to Combined APOBEC3F and APOBEC3G Host Restriction

**DOI:** 10.1101/2024.01.25.577241

**Authors:** Amit Gaba, Maria Yousefi, Shreoshri Bhattacharjee, Linda Chelico

## Abstract

Several APOBEC3 enzymes restrict HIV-1 replication by deaminating cytosine to form uracil in single-stranded proviral (-)DNA. However, HIV-1 Vif binds to APOBEC3 enzymes and counteracts their activity by inducing their proteosomal degradation. This counteraction by Vif is not complete as evidenced by footprints of APOBEC3-mediated mutations within integrated proviral genomes of people living with HIV-1. The APOBEC3 enzymes are co-ordinately expressed in CD4^+^T cells and relative contributions of APOBEC3s in HIV-1 restriction is not fully understood. In this study, we investigated the activity of co-expressed APOBEC3F and APOBEC3G against HIV-1 Subtype B and Subtype C Transmitted/Founder viruses. APOBEC3F and APOBEC3G when co-expressed were previously determined to form a hetero-oligomer that enables partial resistance of APOBEC3F to Vif-mediated degradation. Here, we determined that that APOBEC3F interacts with APOBEC3G through its N-terminal domain. We provide evidence that this results in protection from Vif-mediated degradation because the APOBEC3F N-terminal domain contains residues required for recognition by Vif. We also found subtype specific differences in activity of Transmitted/Founder Vifs against APOBEC3G and the APOBEC3F/APOBEC3G hetero-oligomer. HIV-1 Subtype C Vifs were more active in counteracting APOBEC3G compared to HIV-1 Subtype B Vifs when APOBEC3G was expressed alone. However, HIV-1 Subtype C Vifs were less active against APOBEC3G when APOBEC3F and APOBEC3G were co-expressed. Consequently, when APOBEC3F and APOBEC3G were expressed together HIV-1 Subtype C viruses showed a decrease in relative infectivity compared to that when APOBEC3G was expressed alone. Inspection of Vif amino acid sequences revealed that that differences in amino acids adjacent to conserved sequences influenced the Vif-mediated APOBEC3 degradation ability. Altogether, the data provide a possible mechanism for how combined expression of APOBEC3F and APOBEC3G could contribute to mutagenesis of HIV-1 proviral genomes in the presence of Vif and provide evidence for variability in the Vif-mediated degradation ability of Transmitted/Founder viruses.

**Author Summary:** APOBEC3 enzymes act as barriers to HIV infection by inducing cytosine deamination in proviral DNA, but their effectiveness is hindered by their counteraction by HIV Vif, which leads to APOBEC3 proteasomal degradation. The APOBEC3-Vif interaction has largely been determined using lab adapted HIV-1 Subtype B viruses and with singular APOBEC3 enzymes. Here we examined how primary isolates of HIV-1 replicated in the presence of APOBEC3F and APOBEC3G. APOBEC3F and APOBEC3G interact and this imparts partial resistance to Vif-mediated degradation. We determined that APOBEC3F interacts with APOBEC3G through its N terminal domain, and that APOBEC3F, like APOBEC3G has Vif-mediated degradation determinants in the N-terminal domain, providing a rational for protection from Vif-mediated degradation. We also demonstrate subtype-specific differences in the activity of Transmitted/Founder Vifs against APOBEC3G and the APOBEC3F/APOBEC3G hetero-oligomer. Through an analysis of Vif amino acid sequences, we identified variations influencing the Vif-mediated APOBEC3 degradation ability. This research uncovers previously unidentified mechanisms by which combined expression of APOBEC3F and APOBEC3G may contribute to HIV-1 proviral genome mutagenesis in the presence of Vif and emphasizes the contribution of amino acid variation outside of previously identified conserved regions in Vif-mediating APOBEC3 degradation.

## Introduction

The APOBEC3 (A3) enzymes belong to a family of cytosine deaminases that in primates and humans has at least seven members (A3A-A3H, excluding E) (1). These enzymes act as host restriction factors against retroelements, retroviruses, and DNA viruses (2–4). APOBEC3 enzymes deaminate cytosine to uracil in single-stranded (ss) DNA replication intermediates, which induces mutations in or degradation of DNA, as well as inhibiting retroelement and retroviral reverse transcriptase through a deamination-independent mechanism (2–6).

Out of the seven human A3 family members, five (A3C S188I, A3D, A3F, A3G and A3H (Haplotypes II, V, and VII)) have been reported to inhibit HIV-1 in absence of its Vif protein (7–12). Vif induces degradation of A3 enzymes through the proteasome (13). In the absence of HIV Vif, APOBEC3 enzymes can become encapsidated into budding virions by binding RNAs that are also bound by HIV Gag, such as HIV genomic RNA (14, 15). When these virions infect new cells, A3 enzymes will deaminate cytosines to uracils on HIV single-stranded (-)DNA, resulting in G-to-A mutations in the (+) DNA and the creation of a non-functional provirus (4). Vif acts to prevent this restriction activity by acting as the substrate adaptor in a Cullin RING ligase-5 (CRL5) E3 ligase complex, which binds A3 enzymes through Vif and results in their polyubiquitination and degradation through the 26S proteasome (13). Specifically, HIV-1 Vif becomes complexed with the scaffold protein Cullin 5, Elongin C, and the co-transcription factor CBF-β (13). The Cullin 5 binds Ring finger protein Rbx2 and Elongin C is an obligate dimer with Elongin B (13). This results in a six member complex to which an A3 and ARIH2 or an E2 associates to carry out initial monoubiquitination and polyubiqitination, respectively (16). Vif requires CBF-β not directly for A3-mediated ubiquitination, but for thermodynamic stability in cells, which also facilitates assembly with the CRL5 E3 ligase complex (17–19). However, counteraction of APOBEC3 by Vif is not complete and footprints of A3 induced mutations can be found in at least 30% of integrated proviral genomes within 6 weeks of infection (20–25). At this early stage of infection the viruses are categorized as transmitted/founder (T/F) viruses (26). Since the Vifs of T/F viruses have not been previously characterized for their ability to suppress A3-mediated restriction, the mechanism behind accumulation of this high number of G→A mutations is not understood.

Although there are five human A3 enzymes that can restrict HIV-1, due to the *A3* genes being highly polymorphic, humans usually do not express five *A3* genes that result in enzymes able to restrict HIV (27). The least polymorphic A3 enzymes that have the highest probability to be active against HIV-1 and co-expressed are A3G and A3F (28, 29). Importantly, A3G and A3F form a hetero-oligomer that enhances HIV-1 restriction in the absence of Vif and in the presence of Vif enables the A3F in the hetero-oligomer to become partially resistant to Vif-mediated degradation by a yet-to-be determined mechanism (30, 31). Originally, alanine scanning mutagenesis was used to determine that A3G and A3F interacted with Vif using distinct domains present in the N-terminal and C-terminal domains, respectively (32–35). However, another study using single nucleotide changes based on species specific amino acid differences in A3F determined that the equivalent amino acid at position 128 in the A3G N-terminal domain (NTD) important for Vif-mediated degradation was also important for A3F mediated degradation (36).

Although cryo-EM structures of Vif/CBF-β/ElonginB/ElonginC (VCBC) have been determined for A3G and A3H Haplotype II, there is no structural information for how full length A3F interacts with Vif, thus requiring a dependence on mutagenesis studies to be maintained (37–40). A cryo-EM of the A3F C-terminal domain (CTD) with Vif/CBF-β determined that the Vif and CBF-β interface forms a wedge in which the A3F CTD interacts with both proteins (41). The absence of a structure with full length A3F interacting with Vif has hindered our understanding on how the A3F/A3G hetero-oligomer results in partial resistance of A3F to Vif-mediated degradation.

In this study, we examined both the mechanism by which A3F resists Vif-mediated degradation in the presence of A3G and if this occurs across a panel of HIV-1 T/F viruses. We identified that A3F interacts with A3G through its NTD and confirmed that the NTD of A3F is required for Vif-mediated degradation, suggesting an A3G-induced protection mechanism from ubiquitination. We determined that his occurred not only with HIV-1_LAI_, but also all nine different HIV-1 T/F viruses from Subtypes B and C. There were differences in the capability of T/F Vifs to induce degradation of A3G alone, A3F alone, and the A3F/A3G hetero-oligomer despite previously identified interaction domains being conserved. Rather, we determined that regions adjacent to these conserved domains determined differences in HIV-1 T/F Vif-induced degradation, particularly amino acids at positions 155 and 177. Altogether, our data redefine the Vif interaction domain for A3F and for the first time characterize the Vif-mediated degradation of A3G and A3F from HIV-1 T/F viruses.

## Results

### A3G protects A3F from HIV-1LAI Vif-mediated degradation

A3G and A3F interact through a protein-protein interaction in the absence of an RNA intermediate and this results in partial protection of A3F from Vif-mediated degradation, however the mechanism is not known (30, 31). We studied the A3F/A3G hetero-oligomer using a transfection-based system and a plasmid with two MCS so that each cell that expresses A3F, will also express A3G. The A3F and A3G are uniquely tagged with V5 and HA, respectively, so that they can be identified by immunoblotting. While this precludes comparison between A3G and A3F, we can compare changes among each A3 when it is expressed alone or together to interpret effects of the co-expression.

Samples from HIV-1_LAI_ producer cells transfected with A3F-V5, A3G-HA, or A3F-V5/A3G-HA were immunoblotted to detect steady state protein levels in cell lysates (Fig 1A). Previous findings using VSV-G pseudotyped virus were confirmed with a full-length molecular clone of HIV-1_LAI_. That Vif is responsible for this degradation is demonstrated by the HIV-1_LAI_ SLQ^M^ that has the Vif SLQ motif needed to interact with Elongin C mutated to AAA (Fig 1A). The HIV-1_LAI_ SLQ^M^ condition represents the total possible amount of steady state A3 levels in cells (Fig 1A, Cell). This molecular clone expresses Vif, but it cannot form a CRL5 E3 ubiquitin ligase complex. The data demonstrated that Vif-induced degradation by HIV-1_LAI_ of A3F occurs when it is expressed alone (by comparison to HIV-1_LAI_ SLQ^M^), but when it is co-expressed with A3G, the A3F is more resistant to Vif-mediated degradation (Fig 1A, Cell).

**Fig. 1.**
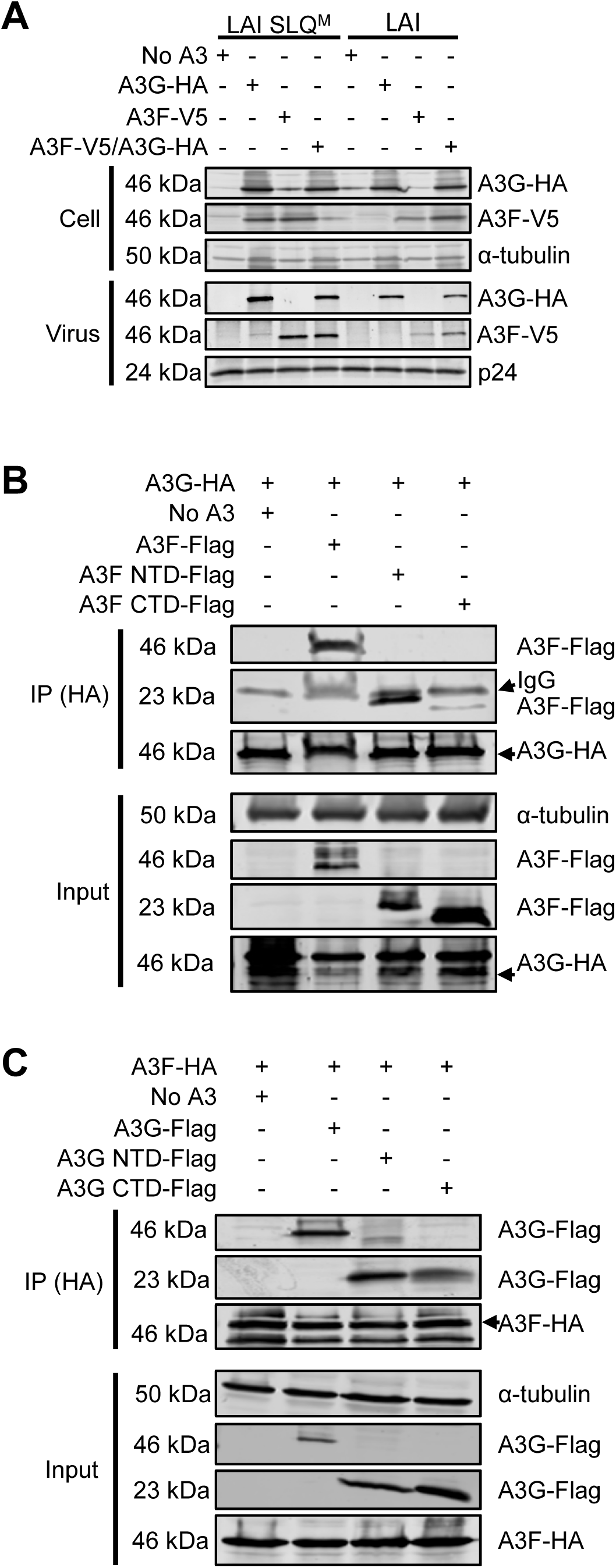
The A3F and A3G interaction results in partial protection of A3F from HIV-1_LAI_ Vif-mediated degradation. **(A)** HIV-1_LAI_ (LAI) and LAI SLQ mutant (SLQ^M^) viruses were used to determine the Vif-mediated degradation of A3F and A3G. The SLQ^M^ is SLQ→AAA and prevents Vif from interacting with Elongin C. Immunoblotting with anti-HA and anti-V5 was used to detect A3G and A3F respectively, in cell and virus lysates when each was expressed alone (A3G-HA, A3F-V5) or co-expressed (A3F-V5/A3G-HA). The anti-α-tubulin and anti-p24 were used as loading controls for cell and virus lysates, respectively. **(B)** Cell lysates co-transfected with expression plasmids for A3G-HA and A3F-Flag, A3F NTD-Flag, or A3F CTD-Flag were immunoprecipitated with anti-HA antibody, resolved with SDS-PAGE and transferred to nitrocellulose membrane. Immunoblotting with anti-HA and anti-V5 demonstrated that the A3F NTD interacted with A3G. The anti-α-tubulin was used as loading control for the input. **(C)** Cell lysates co-transfected with expression plasmids for A3F-HA and A3G-Flag, A3G NTD-Flag, or A3G CTD-Flag were immunoprecipitated with anti-HA antibody, resolved with SDS-PAGE and transferred to nitrocellulose membrane. Immunoblotting with anti-HA and anti-V5 demonstrated that predominantly the A3G NTD, but also the A3G CTD interacted with A3F. The anti-α-tubulin was used as loading control for the input.

The differences in the levels of A3F when co-expressed with A3G compared to when expressed alone was also apparent in HIV-1_LAI_ virions. The additional A3F in cells in the A3F/A3G condition resulted in increased A3F encapsidation into virions (Fig 1A, Virus). However, the amount of A3G encapsidated in virions was similar in the absence or presence of A3F (Fig 1A, Virus). Altogether, these data demonstrate that the increased steady state level of A3F in cell lysates where A3G is also present results in increased encapsidation of A3F and is due to protection from Vif-mediated degradation.

### A3F interacts with A3G through the N-terminal domain

Since the A3F/A3G hetero-oligomer results in protection of A3F from Vif-mediated degradation, but not A3G, we hypothesized that the A3F/A3G interface occludes the residues of A3F that interact with Vif. To determine where A3F and A3G interacted and if this overlapped with the A3F and Vif interface, we used domain constructs of A3F and A3G. A3F and A3G are double zinc deaminase domain (ZDD) enzymes and the two ZDD domains can be stably expressed independently since they are connected by a linker region (42). We used co-IP determine if full length A3G-HA or A3F-HA interacted with Flag tagged constructs of the NTD or CTD of A3F or A3G, respectively. The A3G-HA resulted in strong immunoprecipitation of full length A3F and the A3F NTD, but not the A3F CTD (Fig 1B). These data demonstrated that the A3F NTD interacts with A3G. Similarly, to identify the region of A3G that interacts with A3F we used A3F-HA and A3G full length, NTD, and CTD -Flag constructs. A3F-HA resulted in immunoprecipitation of full length A3G and primarily the A3G NTD, but also a lesser amount of the A3G CTD (Fig 1C). These data support a model in which the A3F NTD is primarily interacting with the A3G NTD. While A3G NTD has been shown to interact with Vif, current structural and mutation data with A3F support that the CTD interacts with Vif (33, 41). As a result, these data do not fully explain how A3G can protect A3F from Vif-mediated degradation.

### A3F NTD is a determinant in Vif-mediated degradation

Although previous structural and biochemical studies have reported that the CTD of A3F interacts with Vif and CBF-β, these data resulted from alanine scanning mutagenesis or Cryo-EM of only the A3F CTD (33, 41). However, even the original study showed that an A3F NTD-A3G CTD chimera interacted with Vif through the A3F NTD and structural models show that the A3F NTD may interact with Vif α1 helix (33, 43). In addition, a single point mutation at A3F amino acid 128, the equivalent amino acid that is a major determinant in the A3G and Vif interaction, can disrupt Vif-mediated degradation of A3F (36). Collectively, these data suggest a previously unidentified key determinant in the A3F NTD for Vif-mediated degradation. As a result, the A3F NTD-A3G NTD interaction may result in occlusion of the A3F residues needed for Vif to efficiently induce A3F degradation, resulting in partial resistance.

To confirm and extend past studies, we used the A3F R128T mutant that showed increased restriction compared to A3F of HIV-1 T/F CH077 (36). We compared the HIV-1_LAI_ and HIV-1 T/F CH077 Vif-mediated degradation of A3F and A3F R128T (Fig 2). We measured the steady state levels of A3F and A3F R128T in producer cells of HIV-1_LAI_, HIV-1_LAI_ SLQ^M^, HIV-1 T/F CH077, and HIV-1 T/F CH077 -Vif. For both HIV-1_LAI_ and HIV-1 T/F CH077 the data showed that A3F R128T is more resistant to Vif-mediated degradation than A3F (Fig 2A). That Vif is responsible for the lowered steady state levels of the A3F is demonstrated by the LAI SLQ^M^ and TF077 -Vif producer cells that do not affect the A3F or A3F R128T steady state levels in cells (Fig 2A).

**Fig. 2.**
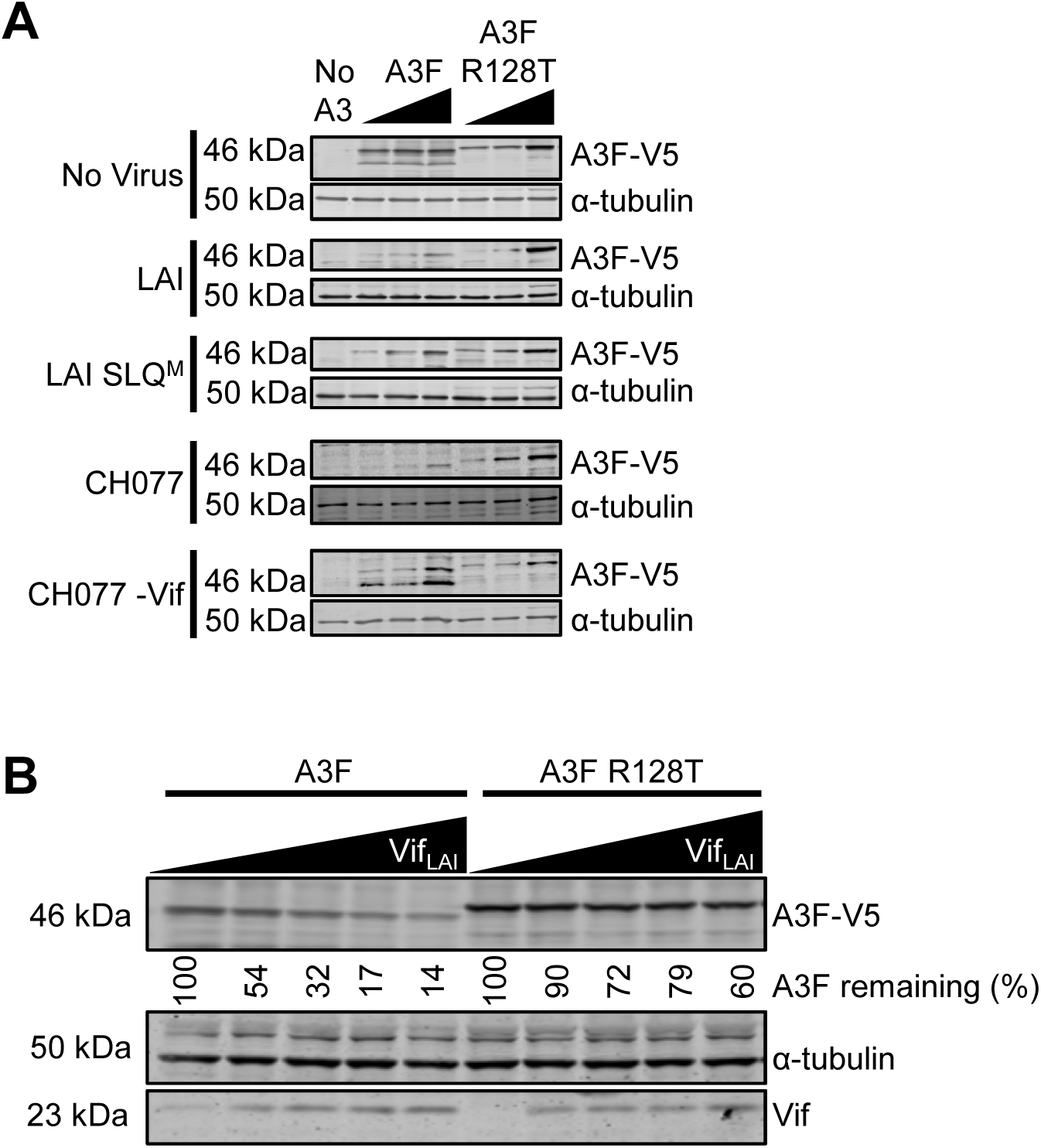
The A3F NTD is a determinant for Vif-mediated degradation. **(A)** Cells were transfected with molecular clones of HIV-1_LAI_, HIV-1_LAI_ SLQ^M^, HIV-1 T/F CH077, or HIV-1 CH077 -Vif and increasing amounts of A3F-V5 (50, 100, or 200 ng) or A3F-V5 R128T (100, 200, or 300 ng) expression plasmid. Cell lysates were collected at 48 h, resolved by SDS-PAGE, transferred to nitrocellulose membrane, and probed with anti-V5 and anti-α-tubulin antibodies. The α-tubulin served as the loading control. **(B)** Cells were transfected with a constant amount of A3F-V5 (100 ng) or A3F-V5 R128T (200 ng) expression plasmid and increasing amounts of HIV-1_LAI_ Vif expression construct (50, 100, 200, or 300ng). Cell lysates were collected at 48 h, resolved by SDS-PAGE, transferred to nitrocellulose membrane, and probed with anti-V5 and anti-Vif antibodies with α-tubulin serving as loading control. The blot was analyzed to determine the band intensity of the A3 with each normalized to the α-tubulin loading control in the same sample. The values were converted to A3F remaining by normalizing to the 0 ng of transfected Vif_LAI_ condition, which was set to a value of 100%.

We also conducted an experiment with a Vif expression plasmid and transfected increasing amounts into cells with a constant amount of A3F or A3F R128T. With increasing amounts of Vif transfected, there was increased degradation of A3F and A3F R128T, however at maximum Vif amounts, the A3F remaining was only 14% but for A3F R128T it was 60% (Fig 2B). At the lowest Vif transfection condition, the A3F R128T was almost completely resistant to Vif-mediated degradation, but for the A3F, there was a 2-fold decrease in the steady state protein level (Fig 2B). Thus, the A3F R128T mutant is more resistant to Vif-mediated degradation than A3F. Taken together this data demonstrate that the A3F NTD is important for Vif-mediated degradation of A3F and amino acid 128 is a key residue. As a result, the A3G likely occludes this region of A3F, making the Vif-mediated degradation less efficient.

### A3G partially protects A3F from HIV-1 T/F Vif-mediated degradation

To determine if the co-expression of A3F and A3G resulted in partial protection of A3F from Vif expressed from primary isolates, we tested nine different HIV-1 T/F viruses from Subtype B (CH058, CH470, CH040, CH077, Thro) and Subtype C (CH236, CH850, CH569) alongside HIV-1_LAI_. Analysis of amino acid sequences from HIV-1 T/F Vifs revealed on average 12% variation relative to Vif of HIV-1_LAI_ (Fig 3A). The maximum variation was 19% for TF236 and the minimum variation was 7% for T/F Thro (Fig 3A). Further inspection of the sequences revealed that the amino acid sequence of the domains reported to be relevant for A3-Vif interactions were conserved and only intervening regions contained the variable amino acids (Fig 3B). Thus, we hypothesized that all the T/F Vifs of Subtype B or Subtype C would have similar degradation profiles for A3G, A3F and A3F/A3G when compared to LAI Vif.

**Fig. 3.**
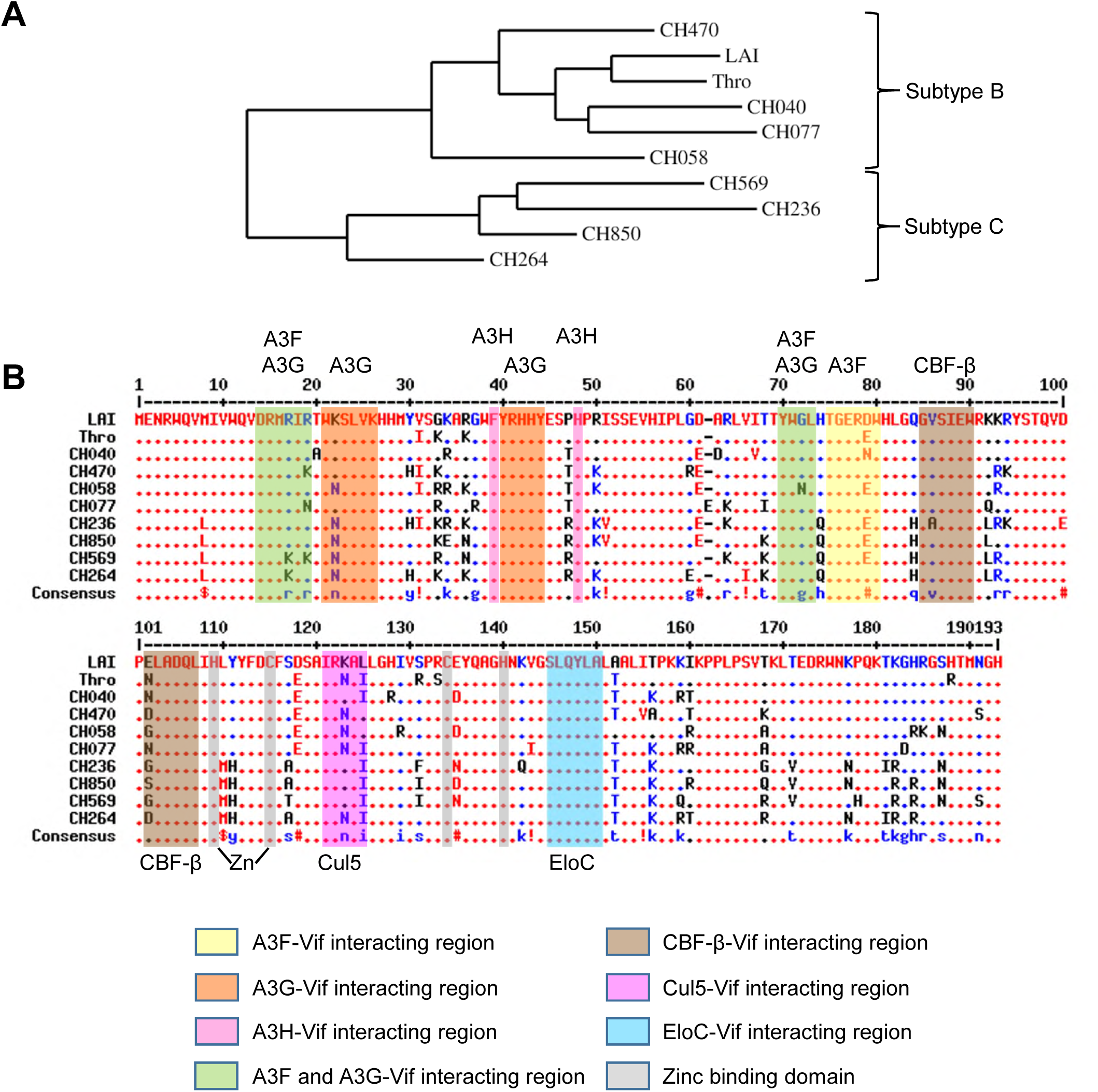
Phylogeny and amino acid alignment of HIV-1 T/F Vifs. **(A)** Phylogenetic analysis of Vif amino acid sequences of TF viruses used in this study was done using the web server phylogeny.fr (63). **(B)** Alignment of Vif amino acid sequences from HIV-1_LAI_ and HIV-1 T/F viruses used in this study. HIV-1_LAI_ was used as the consensus Vif sequence (top). Dots indicate exact amino acid identity, amino acids indicate differences, and dashes indicate gaps introduced to optimize the alignment. Red indicates high consensus, blue indicates low consensus, and black is neutral. The symbols in the consensus sequence indicate the following amino acids: != I, V; $ =L, M; and #=B, D, E, N, Q, Z where B is D or N and Z is E or Q. Known Vif interaction motifs are indicated and were obtained from (55, 64–78). EloC is Elongin C, Cul5 is Cullin 5, Zn is Zinc binding domain. Alignment was done with MultAlin (79).

We selected three HIV-1 T/F molecular clones to study in more depth using single-cycle infectivity assays. The HIV-1 T/F CH040 was used alongside HIV-1_LAI_ to represent Subtype B viruses. The HIV-1 T/F CH850 and CH569 were used to represent Subtype C viruses. We transfected either 25 or 50 ng of plasmids expressing the A3G alone, A3F alone, or A3F/A3G together with the full length molecular clones and determined the relative infectivity in comparison to a no A3 transfection and detected the A3s in producer cells and virions by immunoblotting (Fig 4). The HIV-1_LAI_ SLQ^M^ was used for comparison in each experiment to determine the amount of A3 enzymes in producer cells and viruses in the absence of Vif-mediated degradation and as an experimental control for maximum A3 restriction (Fig 4 and Fig S1).

**Fig. 4.**
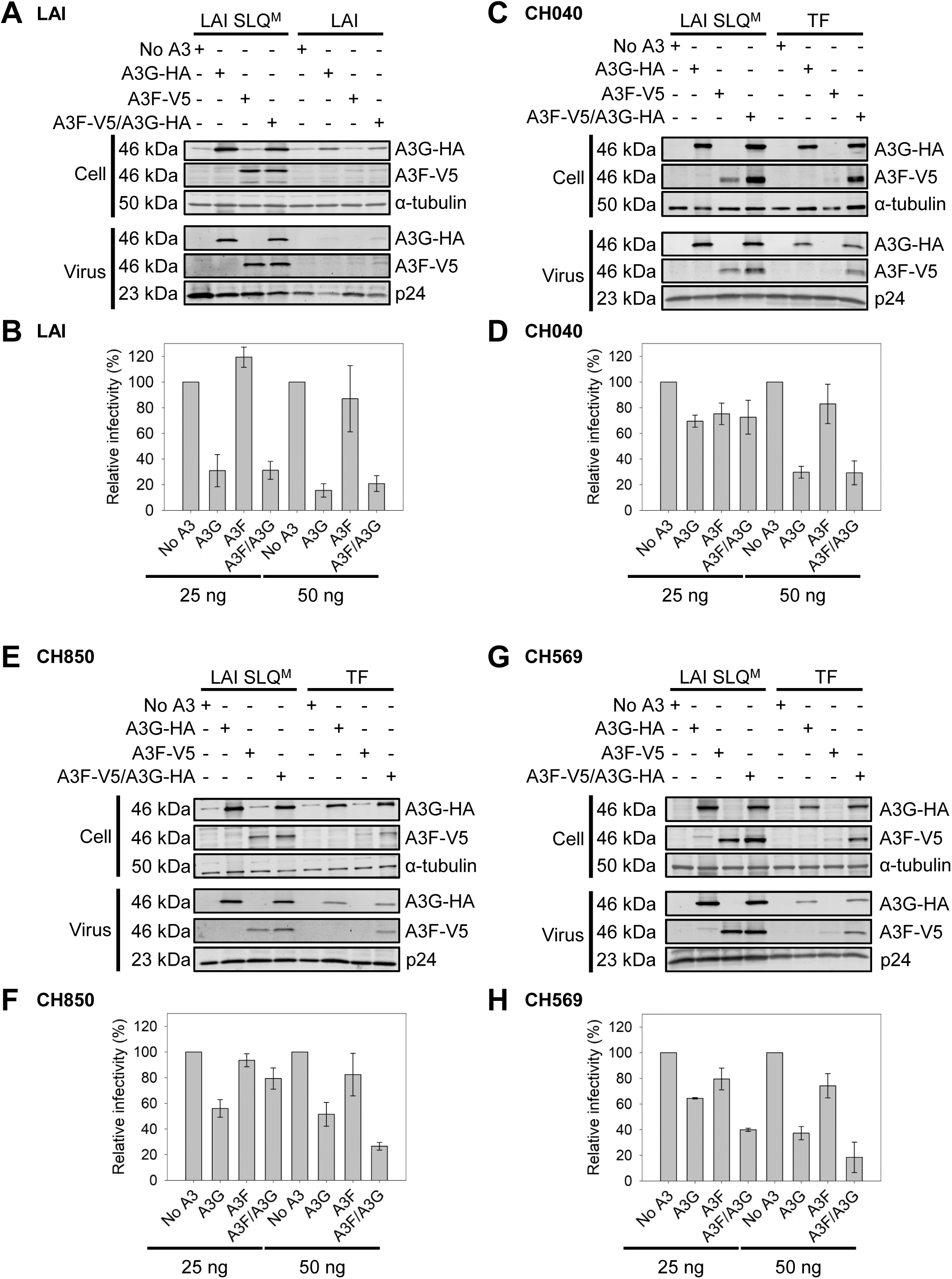
HIV-1 T/F viruses have variable infectivity in the presence of A3F, A3G, and A3F/A3G. 293T cells were co-transfected with No A3 (empty vector), A3G-HA (A3G), A3F-V5 (A3F) or A3F-V5/ A3G-HA (A3F/A3G) expression plasmid and a molecular clone of **(A-B)** HIV-1_LAI_ **(C-D)** HIV-1 T/F CH040 **(E-F)** HIV-1 T/F CH850 or **(G-H)** HIV-1 T/F CH569. For immunoblots, cell lysates and virus lysates were collected at 48 h resolved by SDS-PAGE and transferred to nitrocellulose membrane for probing with anti-V5 and anti-HA. The HIV-1_LAI_ SLQ^M^ was used as a no Vif control. The anti-α-tubulin and anti-p24 served as loading control for cell and virus lysate, respectively. For immunoblots, 50 ng of A3 expression plasmid was used. The relative infectivity was determined using relative β-galactosidase activity from infected TZM-bl cells normalized to the No A3 condition. Viruses used to infect TZM-bl cells were produced from cells transfected with 25 or 50 ng of A3 expression plasmid. Error bars represent the error range from two independent experiments.

Although we observed protection of A3F by A3G in the presence of HIV-1_LAI_ by immunoblotting (Figs 1A and 4A), the co-expressed A3F and A3G were not able to restrict HIV-1_LAI_ more than A3G alone, similar to what was reported previously (Fig 4B) (31). The A3G remaining in the cells when alone or co-expressed with A3F was encapsidated (Fig 4A, Virus). This was in contrast to A3F that was not detectable in virions by immunoblotting when expressed alone, but was able to be detected in virions when co-expressed with A3G (Figs 1A and 4A, Virus). However, when determining the infectivity, we observed that HIV-1_LAI_ was restricted similarly when A3G was expressed alone and when expressed with A3F (Fig 4B). Mohammadzadeh *et al*., found similar results for infectivity, but also determined that there was an effect of the A3F/A3G hetero-oligomer upon examining the mutations in proviral sequences (31). The HIV-1_LAI_ was sensitive to restriction when A3G was expressed, despite the presence of Vif and at both 25 and 50 ng transfection conditions. HIV-1_LAI_ was not sensitive to A3F-mediated restriction, consistent with previously reported lower activity of A3F and the increased sensitivity to Vif-mediated degradation (Fig 4A-B) (29, 31). However, the HIV-1 T/F viruses showed different infectivity results from HIV-1_LAI_, although the immunoblots were similar.

Specifically, immunoblotting of producer cell and virus lysates from Subtype B HIV-1 T/F CH040 single-cycle infectivity experiments showed modest A3G degradation when alone or co-expressed with A3F and efficient A3F degradation when alone, but not when co-expressed with A3G (Fig 4C, Cell). The A3s that were observed in the cell lysates (A3G alone, A3G co-expressed with A3F, and A3F co-expressed with A3G) were encapsidated (Fig 4C). Despite there being a significant protection of A3F from Vif-mediated degradation and encapsidation of A3F in the presence of A3G, the infectivity data showed no effect of A3G alone, A3F alone, or A3F/A3G on T/F CH040 when only 25 ng of the A3s was transfected (Fig 4C-D). When the transfection level of the A3s was doubled, we observed A3G, but not A3F restriction when both were expressed alone the effect of A3F/A3G co-expression was similar to A3G alone (Fig 4D).

The HIV-1 Subtype C T/F viruses CH850 and CH569 showed different infectivity results from the Subtype B viruses. Interestingly, the immunoblots for both HIV-1 T/F CH850 and CH569 were similar to those of the Subtype B viruses and showed modest A3G degradation when alone or co-expressed with A3F, efficient degradation of A3F when expressed alone, and protection of A3F from Vif-mediated degradation when co-expressed with A3G (Fig 4E, G). For HIV-1 T/F CH850, the infectivity was not affected by expression of A3G alone, A3F alone or A3F/A3G at 25 ng plasmid transfection. However, at the 50 ng plasmid transfection we observed no difference in the sensitivity to A3G or A3F alone, but there was ∼2-fold more sensitivity to A3F/A3G compared to A3G alone, resulting in only ∼30% remaining infectivity (Fig 4F). There was even more of an effect of co-expression of A3F/A3G for HIV-1 T/F CH569 where we observed ∼2-fold more reduction in infectivity for A3F/A3G compared to A3G alone at both the 25 and 50 ng plasmid transfections with only ∼20% infectivity remaining at the higher A3F/A3G transfection amount (Fig 4H). These effects are noteworthy considering that Vif expression is occurring from these molecular clones.

Overall, we observed similar results for other Subtype B and C HIV-1 molecular clones (S1 Fig). The Subtype B molecular clones were more sensitive to restriction by A3G when expressed alone and showed no difference from A3G alone for the restriction in the A3G/A3F condition (S1 Fig). The Subtype B molecular clones also showed a large variability in sensitivity to A3G and A3F/A3G, with infectivity ranging from 15-54% (S1 Fig). The Subtype C molecular clones were less sensitive to restriction by A3G when expressed alone, but showed an approximately 2-fold reduction in infectivity in the presence of A3F/A3G in comparison to A3G alone (S1 Fig). None of the molecular clones were sensitive to restriction by A3F when it was expressed alone (S1 Fig).

### HIV-1 T/F Subtype C specific differences in A3G-mediated degradation

Despite clear trends emerging from the infectivity data and corresponding immunoblots, we wanted to conduct a more quantitative assay to characterize the potential differences in the Vif-mediated degradation of Subtype B and Subtype C T/F virus Vifs. To this end, we cloned the *vif* from HIV-1_LAI_ and the T/F viruses CH040, CH850, and CH569. We used these clones for A3 degradation assays using co-transfection of a constant amount of A3G, A3F or A3F/A3G expression plasmid and increasing amounts of Vif expression plasmid. For the Subtype B Vifs from HIV-1_LAI_ and T/F CH040, there was less A3G remaining in the A3F/A3G condition in comparison to Subtype C virus Vifs (Fig 5A, C, E, G and Table 1, 25 ng). Specifically, for T/F CH040 there was 2-fold to 3-fold less A3G in the A3F/A3G condition when compared to CH850 and CH569, respectively (Fig 5C, E, G and Table 1, 25 ng Vif plasmid transfected). This suggests that HIV-1 Subtype B Vifs are more efficient than Subtype C viruses in causing degradation of A3G when it is co-expressed with A3F. There was no consistent subtype differences in the ability of Vif to induce degradation of A3G when expressed alone.

**Fig. 5.**
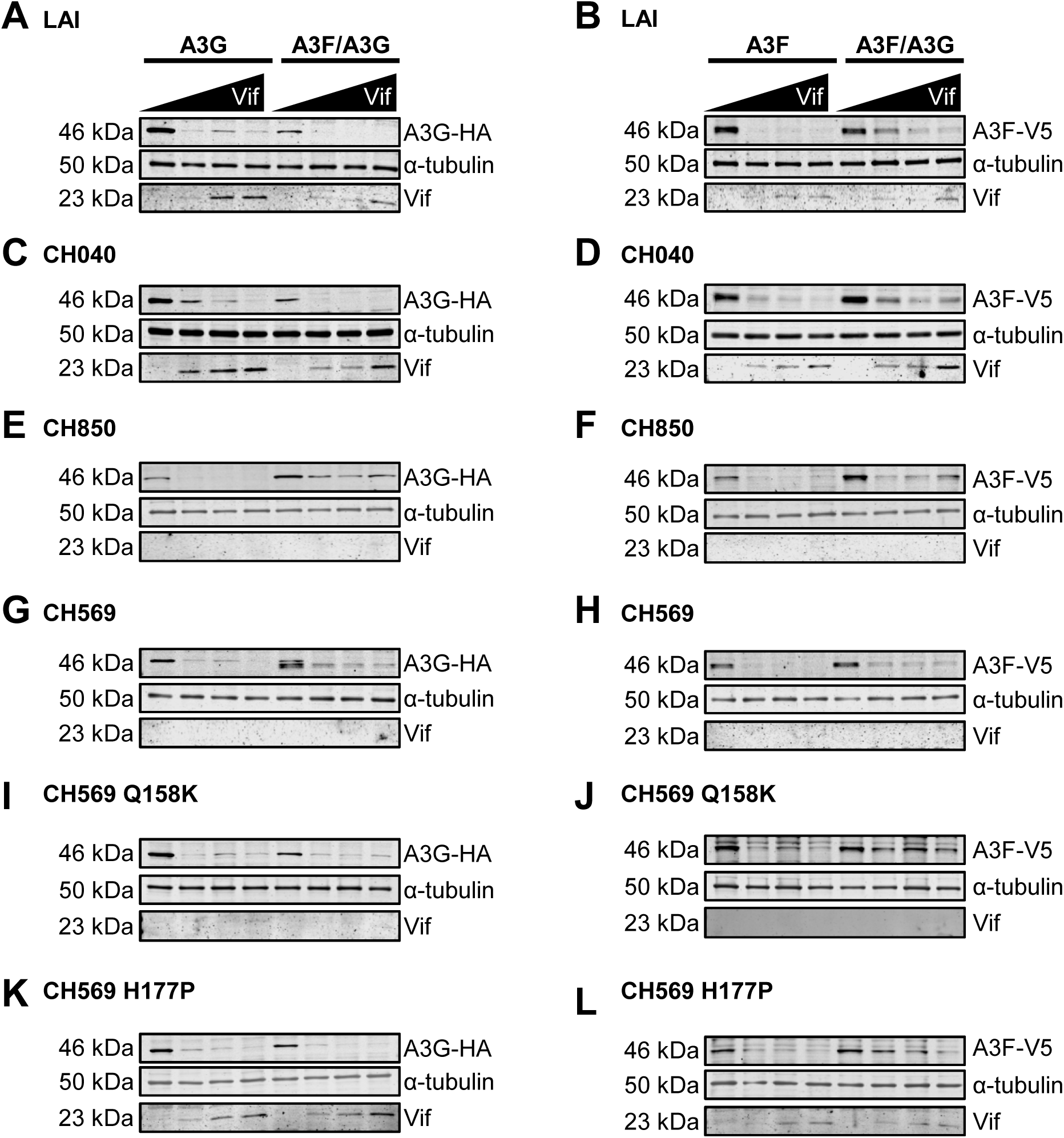
Vif-mediated degradation of A3F, A3G, and A3F/A3G differs between HIV-1 Subtype B and Subtype C viruses. 293T cells were co-transfected with plasmid expressing either A3G-HA (A3G), A3F-V5 (A3F) or A3F-V5/A3G-HA (A3F/A3G) and increasing amount of Vif-AU1 expression plasmid from HIV-1 **(A-B)** LAI **(C-D)** T/F CH040 **(E-F)** T/F CH850, **(G-H)** T/F CH569, **(I-J)** T/F CH569 Q158K, and **(K-L)** T/F CH569 H177P. Cell lysates were collected at 44 h, resolved by SDS-PAGE and transferred to nitrocellulose membrane for probing with anti-V5, anti-HA and anti-Vif antibody. The anti-α-tubulin was the loading control.

**Table 1.**
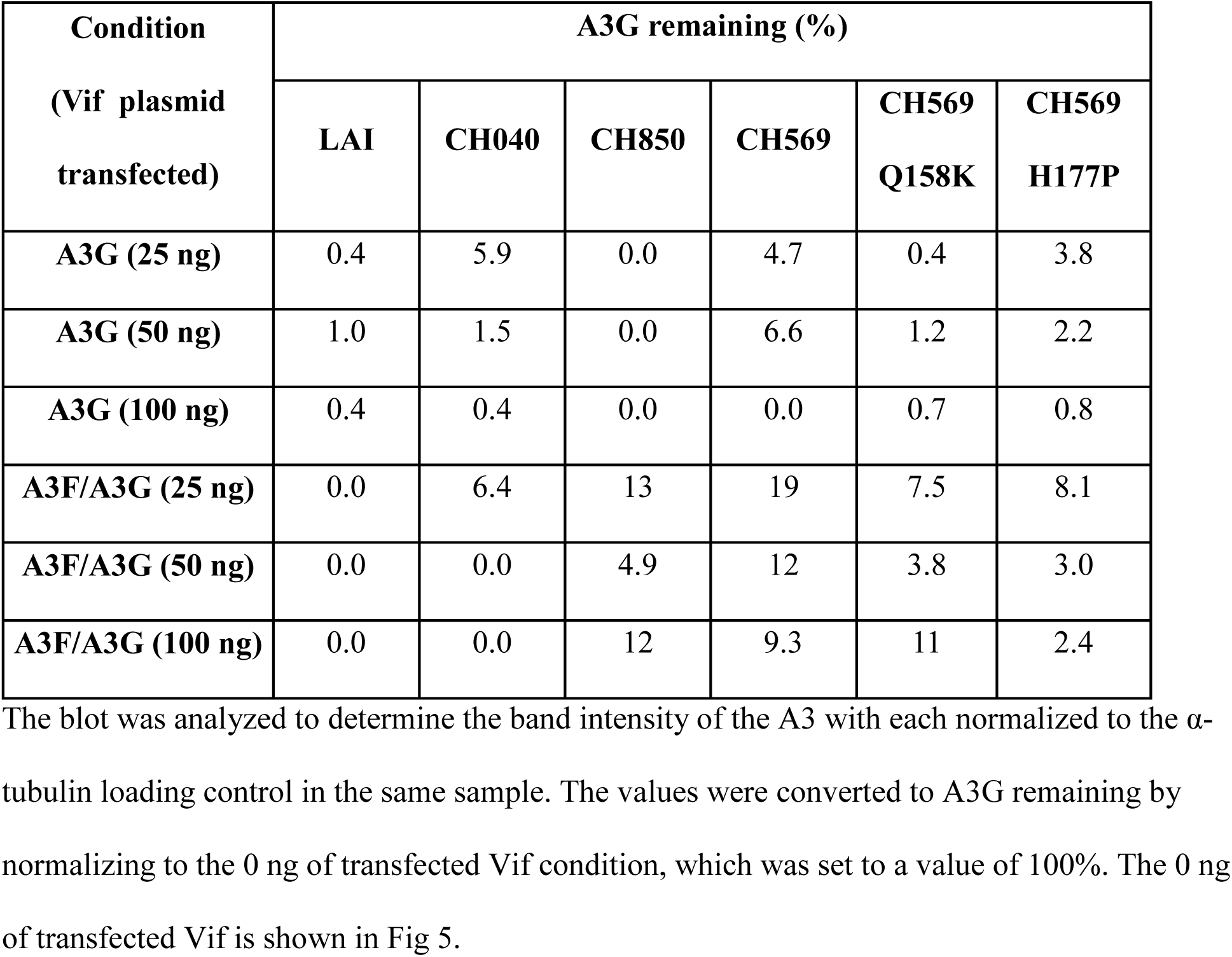
Relative A3G detected by immunoblotting in the presence of Vif.

| **Condition**<br>**(Vif plasmid**<br>**transfected)** | **A3G remaining (%)** |  |  |  |  |  |
| --- | --- | --- | --- | --- | --- | --- |
|  | **LAI** | **CH040** | **CH850** | **CH569** | **CH569**<br>**Q158K** | **CH569**<br>**H177P** |
| **A3G (25 ng)** | 0.4 | 5.9 | 0.0 | 4.7 | 0.4 | 3.8 |
| **A3G (50 ng)** | 1.0 | 1.5 | 0.0 | 6.6 | 1.2 | 2.2 |
| **A3G (100 ng)** | 0.4 | 0.4 | 0.0 | 0.0 | 0.7 | 0.8 |
| **A3F/A3G (25 ng)** | 0.0 | 6.4 | 13 | 19 | 7.5 | 8.1 |
| **A3F/A3G (50 ng)** | 0.0 | 0.0 | 4.9 | 12 | 3.8 | 3.0 |
| **A3F/A3G (100 ng)** | 0.0 | 0.0 | 12 | 9.3 | 11 | 2.4 |

An additional difference was that for HIV-1 Subtype B Vifs more A3F remained during A3F/A3G co-expression compared to Subtype C virus Vifs (Fig 5B, D, F, H and Table 2). Specifically, for T/F CH040 there was 2-fold more A3F in the A3F/A3G condition than both CH850 and CH569 (Fig 5D and Table 1, 25 ng Vif plasmid transfected). Higher transfection amounts of the Vif plasmid resulted in equivalent amounts of degradation due to saturation of experimental system with Vif (Fig 5D and Table 1). These data suggest that Subtype C virus Vifs are more active in causing degradation of A3F in the A3F/A3G hetero-oligomer compared to Subtype B virus Vifs. However, for T/F CH850 and CH569 there was still some protection of A3F within the A3F/A3G hetero-oligomer, this was approximately 2-fold less that what was observed with HIV-1_LAI_ and T/F CH040. Interestingly, the Subtype C virus Vifs were not detectable on the blots and were presumably degraded with the A3, in contrast to the Subtype B Vifs that could be detected by immunoblotting (Fig 5). Altogether, these data suggest that the HIV-1 T/F Subtype C viruses are more sensitive to the A3F/A3G hetero-oligomer than HIV-1 T/F Subtype B because the Vifs are less able to induce degradation of A3G (Fig 5A, C, E, G).

**Table 2.** Relative A3F detected by immunoblotting in the presence of Vif.

| **Condition**<br>**(Vif plasmid**<br>**transfected)** | **A3F Remaining (%)** |  |  |  |  |  |
| --- | --- | --- | --- | --- | --- | --- |
|  | **LAI** | **CH040** | **CH850** | **CH569** | **CH569**<br>**Q158K** | **CH569**<br>**H177P** |
| **A3F (25 ng)** | 0.0 | 17 | 0.0 | 0.0 | 11 | 41 |
| **A3F (50 ng)** | 0.0 | 6.1 | 0.0 | 0.0 | 16 | 12 |
| **A3F (100 ng)** | 0.0 | 0.0 | 0.0 | 0.0 | 8.1 | 7.0 |
| **A3F/A3G (25 ng)** | 31 | 29 | 12 | 14 | 34 | 30 |
| **A3F/A3G (50 ng)** | 22 | 14 | 14 | 10 | 31 | 20 |
| **A3F/A3G (100 ng)** | 12 | 17 | 24 | 10 | 23 | 9.3 |

### Amino acid determinants for Vif-mediated degradation occur outside of conserved domains

The Vif domains for interaction with CBF-β, Elongin C and Cullin 5 are well conserved since deviation from these motifs would disrupt function. Nonetheless, we examined the amino acid alignment of the HIV-1 T/F Vifs to determine if we could identify any amino acids in Subtype C in comparison to Subtype B viruses that account for differential abilities of the Vifs to induce A3 degradation (Fig 3B). All analyzed Subtype C viruses, but none of the Subtype B viruses had a Leucine at position 8 that is near the DRMR A3F/A3G interaction domain, a Histidine at position 83 and Leucine at position 91 near the CBF-β interaction domain, and a Methionine and Histidine at 110-111 near the CBF-β and Zn^2+^ interaction domains (Fig 3B).

While all these amino acids likely contribute individually or collectively to the differences we observed with infectivity and A3-induced degradation, we focused on CH569 since it had a unique profile for infectivity (Fig 4). The CH569 was most sensitive to the A3F/A3G co-expression and this appeared to be due to a lesser ability to induce degradation of A3G, in addition to an inability to induce full degradation of A3F in the A3F/A3G hetero-oligomer (Fig 4H and Fig 5G-H). We observed that CH569 Vif had two unique amino acids from all other Subtype B and Subtype C Vifs that we tested (Fig 3B). We mutated these amino acids to become more like the other T/F Vifs and LAI Vif, resulting in CH569-Q158K and CH569-H177P Vifs.

We used these CH569 mutant Vifs in an A3 degradation assay. When A3F was co-expressed with A3G, ∼2-fold more A3F was remaining for both CH569 Vif mutants compared to wild type CH569 Vif suggesting these amino acids are required for efficient degradation of A3F in the A3F/A3G hetero-oligomer (Fig 5G-L and Table 2, 25 and 50 ng Vif plasmid transfected). For CH569-Q158K there was also less degradation of A3F alone (Fig 5G-L and Table 2, 25 and 50 ng Vif plasmid transfected). Similarly, the CH569-H177P compared to wild type CH569 resulted in less degradation of A3F when expressed alone and within the hetero-oligomer (Fig 5G-H, K-L and Table 2, 25 ng Vif plasmid transfected). These mutations were predicted to make CH569 more Subtype B-like and the results align well with our HIV-1 Subtype B data where more A3F was remaining in the A3F/A3G expression condition than Subtype C viruses (Fig 5 and Table 2, 25 ng Vif plasmid transfected).

Interestingly, the CH569 Vif mutants had a different effect on Vif-mediated degradation of A3G alone and A3G in the A3F/A3G condition. While the CH569-H177P mutant had no effect on A3G degradation when expressed alone, the CH569-Q158K mutant was ∼6-10 fold more efficient at Vif-mediated degradation of A3G (Fig 5 and Table 1, 25 and 50 ng Vif plasmid transfected). In the A3F/A3G condition, both the CH569 -H177P and -Q158K mutants were ∼2-3-fold more efficient at inducing degradation of A3G and more similar to Vif-mediated degradation observed for Subtype B viruses, as would be predicted by the amino acid substitutions (Fig 5 and Table 1, 25 and 50 ng Vif plasmid transfected).

Overall, these data suggest that there are still unidentified regions of Vif that are determinants in A3-mediated degradation. This is especially important for understanding the A3F and A3F/A3G interactions with Vif since there is no structural information for full length A3F in complex with Vif.

## Discussion

A3 restriction factors are able to induce mutations in HIV-1 proviral DNA despite Vif-mediated degradation. Near full-length sequencing of HIV-1 proviral DNA within the acute infection period has shown that ∼95% of proviral DNA becomes inactivated within the first 6 weeks of infection and at least 30% of those proviruses have A3-induced G→A mutations, particularly causing stop codons that result in viral inactivation (20, 44). While this initially appeared at odds with a functional Vif protein, A3F and A3G have been shown to hetero-oligomerize and co-encapsidate in order to partially resist Vif-mediated degradation enabling their activity to tip the balance towards viral inactivation (30, 31, 45). We have previously shown that in the presence of Vif and antiretrovirals, A3F/A3G hetero-oligomerization results in partial protection of A3F from Vif-mediated degradation, increased proviral mutations from A3F, and greatly decreased viral fitness compared to the absence of A3s (31). However, how A3F and A3G interacted and if these identified hetero-oligomer features occurred in the presence of HIV-1 T/F Vifs was not known. Here, we determined that the A3F NTD interacts with A3G, confirmed and extended data that suggested the A3F NTD is a determinant in Vif-mediated degradation, determined that multiple HIV-1 T/F viruses lose the ability to fully induce degradation of A3F when it hetero-oligomerizes with A3G, and identified new determinants in Vif-mediated degradation outside of the previously identified domains.

The co-IP data demonstrated that A3F the NTD interacted with A3G. An additional consideration in this interaction is the stoichiometry and RNA binding. Ara *et al*. determined that three molecules of A3F interacted with one molecule of A3G (30). However, the orientation of the A3F interacting domains for homo-oligomerization is not known, making it difficult to model the A3F/A3G interaction and how the interaction is able to protect A3F from Vif-mediated degradation. That the NTD of A3F is involved in Vif-mediated degradation and the interaction with A3G suggests that A3G occludes the Vif interaction site, however, it does not explain why other A3F subunits of the hetero-oligomer are also protected from Vif-mediated degradation. One explanation is that both A3G hetero- and A3F homo-oligomerization block the Vif interaction site. Alternatively, since the A3F protection from Vif-mediated degradation is only partial, this could mean that only the A3F interacting with A3G is protected from Vif mediated degradation, and other A3F molecules in the hetero-oligomer are not. Additionally, A3G may also be protected from Vif-mediated degradation when interacting with A3F since the A3G NTD is the Vif interaction domain (13). We may not observe A3G protection from Vif-mediated degradation when interacting with A3F in bulk cell lysate studies because the stoichiometry of the hetero-oligomer involves only one A3G to three A3F (30). Notably, multiple structural studies have demonstrated that Vif interacts with a monomer of A3G and an RNA molecule (37–39). How the RNA molecule associates with the A3F/A3G hetero-oligomer and if A3F also interacts with Vif and RNA remains to be determined.

The A3F/A3G hetero-oligomer may affect Vif-mediated ubiquitination. Studies that determined where the CRL5 E3 ligase polyubiquitinates lysines on A3G and A3F found different results. Iwatani *et al*. determined that A3G was polyubiquitinated on four key residues and these could be mutated to block Vif-mediated ubiquitination (46). However, Albin *et al*. determined that although A3G was partially resistant to Vif-mediated degradation when the four key residues identified by Iwatani *et al*. were mutated, it did not result in complete resistance to Vif (46, 47). These results and that A3F was found to be polyubiquitinated throughout the protein and modification of specific lysines could not block Vif-mediated degradation of A3F resulted in the conclusion that ubiquitination of A3s by the CRL5 E3 ligase is relatively stochastic (47). As a result, we hypothesize an all or nothing phenomenon where those A3F molecules interacting with A3G are not ubiquitinated due to not directly interacting with Vif rather than a hypothesis of less efficient ubiquitination of A3F resulting in partial protection of A3F from Vif-mediated degradation.

Combined restriction of HIV-1 from A3F and A3G in the presence of Vif has resulted in variable conclusions. An early study found that in the absence of Vif, co-expressed A3G and A3F caused an increase in HIV-1 proviral DNA mutations compared to when each was expressed alone (48). However, effect of co-expression of A3F and A3G on infectivity was only additive (48). Although in the absence of Vif Ara *et al*. found that there was a synergistic effect of A3F and A3G co-expression on HIV-1 proviral mutation rate and restriction, in the presence of Vif Mohammadzadeh *et al*. observed that there was no effect on either of these variables (30, 31). Rather, the effects observed by Mohammadzadeh *et al*. were that the proviral mutation type became more diversified and that the viruses were less fit under selective conditions, such as antiretroviral treatment (31). These studies all used HIV-1 Subtype B viruses and our data are in agreement with these studies showing that despite partial protection of A3F from Vif-mediated degradation, in a single-cycle infectivity assay, there is no effect on HIV-1 infectivity above that of A3G alone. There are two hypotheses for these results with HIV-1 Subtype B viruses in our study. This could be due to the lack of a selective pressure that would reveal defects in fitness or that more replication cycles are needed to see a pronounced effect in the presence of Vif. These hypotheses are not mutually exclusive. Mohammadzadeh *et al.* supports the former hypothesis (31). The latter hypothesis is supported by mouse studies. Krisko *et al*. used a BLT humanized mouse model and concluded that for consistent HIV-1 restriction the combined activity of A3F and A3G was required (49). However, Sato *et al*. that utilized a NOG-hCD34 mouse humanized mouse model found that mutations in HIV-1 proviral DNA were caused by deamination activity of either A3G or A3F suggesting that A3F and A3G act individually, but still concluding that they can restrict HIV-1 in the presence of Vif (50).

Surprisingly, there was a consistent and different observation when A3F and A3G co-expression occurred during infection with HIV-1 T/F Subtype C viruses. The four HIV-1 T/F Subtype C viruses used in this study all had a 2-fold decrease in infectivity during A3F/A3G co-expression compared to A3G alone (50 ng A3 plasmid transfection). When compared to HIV-1 T/F Subtype B viruses, this did not result from only a difference in the level of A3F protection from Vif-mediated degradation, but also a lesser ability to induce degradation of A3G when expressed alone and within the A3F/A3G hetero-oligomer. However, when A3G was expressed alone, the HIV-1 T/F Subtype B and Subtype C viruses showed a large diversity in the A3G-mediated restriction with Subtype B viruses on average being more sensitive than Subtype C viruses to A3G-medated restriction. However, during A3F/A3G co-expression the HIV-1 T/F Subtype C viruses were more sensitive to restriction. These data suggest that the additional A3G encapsidated in the presence of A3F for HIV-1 T/F Subtype C viruses, but not Subtype B viruses causes the more efficient restriction. However, since this does not occur when A3G is expressed alone during HIV-1 T/F Subtype C infection, the data support that there is a contribution of restriction from A3F. This may be due to the increased diversity in proviral DNA mutations as identified by Mohammadzadeh *et al.* (31). Since A3F deaminates the 5’TC motif in (-) DNA, it has a greater number of possible codons that this site overlaps with compared to A3G that deaminates 5’C<u>C</u> (underlined C primarily deaminated) and causes mutations primarily at Glycines (70% of total mutations) (51).

The variability in the HIV-1 T/F virus sensitivity to A3G-mediated restriction was surprising since the A3-Vif interaction domains are highly conserved. However, outside of these regions Vifs display variability, particularly subtype specific amino acid variability (52–54). Although, HIV-1 Subtype C is the most prevalent subtype globally and accounts for more than 40% global infections there are few studies using HIV-1 Subtype C Vif to map Vif-A3 interacting regions, in addition to a lack of A3-Vif studies using Vifs from HIV-1 T/F viruses. Our data indicate that Vif amino acids outside the well-defined Vif-A3 interfaces influence the degradation ability. However, the residues we identified as unique to the HIV-1 T/F CH569 Vif were in a loop region not well defined in cryo-EM or structural studies (37–39, 55). Nonetheless, this region is between the Elongin C interaction site and the PPLP motif of Vif that was previously shown to be important for A3-mediated degradation. As a result, additional structure function studies in this region of Vif are warranted.

The data presented here reveal additional details on the A3F/A3G interaction, demonstrate that HIV-1 T/F viruses have differential sensitivities to A3G and A3F/A3G restriction, and that HIV-1 T/F Subtype C viruses are most sensitive than Subtype B viruses to the A3F/A3G hetero-oligomer mediated restriction. Structural studies defining how A3G and A3H interact with VCBC have revealed new facets of the A3-Vif interactions, but there remains no structural information with full length A3F or an understanding of how VCBC interacts with the A3F/A3G hetero-oligomer (37–41). Altogether, the data emphasize that the Vif-A3 interaction is still not completely defined and additional studies utilizing T/F viruses, especially those of Subtype C origin are warranted.

## Materials and Methods

### Plasmids and Molecular Clones

The expression plasmids pVIVO2 A3G-HA, pVIVO2 A3F-V5 and pVIVO2 A3F-V5/A3G-HA, have been previously described (30). The following reagents were obtained through the NIH HIV Reagent Program, Division of AIDS, NIAID, NIH: Plasmid pcDNA3.1-APOBEC3G-HA Expressing Human APOBEC3G with C-Terminal Triple HA Tag, ARP-9952, contributed by Dr. Warner C. Greene and Human APOBEC3F V5 His Expression Vector, ARP-10100, contributed by Dr. B. Matija Peterlin and Dr. Yong-Hui Zheng. The ARP-10100 APOBEC3F plasmid was subcloned in place of A3G in the ARP-9952 vector background. Flag-tagged constructs in pcDNA3.1 for A3F, A3F-NTD, A3F-CTD, A3G, A3G-NTD, and A3G-CTD were provided by Dr. Xiaojiang S. Chen (University of Southern California). The A3F-V5 R128 has been previously described (36). The expression plasmid pcDNA3.1-Vif_LAI_ has been previously described (56). The pCG-Vif-AU1 plasmids were generated by PCR amplifying Vif sequences from infectious molecular clones and inserted using XbaI and MluI sites.

The infectious molecular clone of HIV-1_LAI_ and the HIV-1_LAI_ SLQ→AAA (SLQ^M^) that contains a mutated Vif that cannot interact with Elongin C or induce degradation of A3s has been previously described (57). The infectious molecular clones for the HIV-1 transmitted founder (T/F) viruses were a gift from Dr. Beatrice Hahn (Perelman School of Medicine, University of Pennsylvania) and have been previously described (58–61). The CH077 -Vif construct has two tandem stop codons inserted in *vif* and was previously described (36).

### Co-immunoprecipitation

The 293T cells were seeded in a T75 cm^2^ flask (1 x 10^6^) and the next day were transfected with plasmid DNA using GeneJuice transfection reagent (EMD Millipore) as per manufacturer’s instructions. Following amounts of plasmid DNA were used: pcDNA3.1 A3F-Flag (4 µg), pcDNA3.1 A3F-NTD-Flag (3 µg), pcDNA3.1 A3F-CTD-Flag (2 µg), pcDNA3.1-A3G-HA (1 µg), pcDNA3.1-A3F-HA (4 µg), pcDNA3.1-A3G-Flag (1 µg), pcDNA3.1-A3G-NTD-Flag (2 µg), and pcDNA3.1-A3G-NTD-Flag (2 µg). Empty pcDNA3.1 was used to equalize the amount of transfected DNA. At 48 h post transfection, the cells were washed with PBS and lysed using IP buffer (50 mM Tris-Cl pH 7.4, 1% Nonidet-P40, 10% glycerol, 150 mM NaCl) supplemented with EDTA-free protease inhibitor (Roche) and clarified by centrifugation. Then 400 µg of cell lysate was added to anti-HA magnetic beads (Sigma) in the presence of RNaseA (50 μg/ml; Roche) and incubated for 2 h with gentle rocking at 4°C. The beads were subsequently washed three times with IP buffer, and the immunoprecipitated proteins were eluted with Laemmli buffer and subjected to SDS-PAGE and immunoblotting with anti-Flag antibody (Cat# F1804, Sigma, 1:1,000), anti-HA antibody (Cat # H6908, Sigma, 1:10,000), and anti-α-tubulin antibody (Cat # PA1-20988, Invitrogen, 1:1,000). Secondary detection was performed using Licor IRDye antibodies produced in goat (IRDye 680-labeled anti-rabbit, 1: 10,000 Cat # 926-68071 and IRDye 800-labeled anti mouse, 1:10,000 Cat # 926-32210).

### A3 degradation assay

To investigate the ability of Vif expressed from infectious molecular clones to induce degradation of A3F and A3F R128T, 293T cells (1 × 10^5^ per well) in 12 well plates were co-transfected with pVIVO2 A3F-V5 (50, 100, and 200 ng) or pcDNA3.1 A3F R128T (100, 200, and 300ng) expression vectors and 500 ng of infectious molecular clones of HIV-1_LAI_, HIV-1_LAI_ SLQ^M^, HIV-1 T/F CH077, or HIV-1 T/F CH077 -Vif. At 16 h post transfection the media was changed. After 44 h post transfection, cells were washed with PBS and lysed using 2× Laemmli buffer.

In order to compare the efficiency of HIV-1_LAI_ Vif in mediating the degradation of A3F and A3F R128T, 293T cells (1 × 10^5^ per well) in 12 well plates were co-transfected with 100 ng pcDNA3.1 A3F-V5 or 200 ng pcDNA3.1 A3F-V5 R128T and an increasing amount of pcDNA3.1 Vif_LAI_ (50, 100, 200, or 300 ng). Then, 16 h post transfection the media was changed. At 44 h post transfection, cells were washed with PBS and lysed using 2× Laemmli buffer.

To compare efficiency of different Vifs in mediating degradation of A3G-HA, A3F-V5, and co-expressed A3F-V5/A3G-HA, 293T cells (1 × 10^5^ per well) in 12 well plates were co-transfected with 50 ng of pVIVO2 expressing No A3, A3G-HA, A3F-V5, or A3F-V5/A3G-HA and titration of pCG-Vif-AU1 (0, 25, 50, and 100 ng) using GeneJuice (Novagen) transfection reagent according to manufacturer’s instructions. To equalize the amount of plasmid DNA transfected, empty pCG vector was used. Then, 16 h post transfection the media was changed. At 44 h post transfection, cells were washed with PBS and lysed using 2× Laemmli buffer.

Protein from each cell lysate (30 µg) was resolved by SDS-PAGE used for immunoblotting on nitrocellulose membrane. Primary detection was performed using mouse monoclonal anti-HA antibody (Cat# H9658, Sigma, 1:10,000), anti-V5 mouse monoclonal antibody (Cat# V8012, Sigma, 1:1000), rabbit anti-Vif antibody (Cat# 809, NIH HIV Reagent Program, contributed by Dr. Bryan Cullen, 1:1000) and anti-α tubulin antibody (rabbit polyclonal, Cat# PA1-20988, Invitrogen, 1:1000) Secondary detection was performed using Licor IRDye antibodies produced in goat (IRDye 680-labeled anti-rabbit 1:10,000 Cat # 926-68071 and IRDye 800-labeled anti mouse 1:10,000 Cat # 926-32210). For quantification, Image Studio was used to detect the pixel intensity of the experimental and loading control bands. Each sample was normalized to its own loading control before comparison to other lanes.

### Single-cycle infectivity assay

Single-cycle infectivity assays were carried out using 1 × 10^5^ 293T cells per well in a 12 well plate. The 293T cells were maintained in DMEM with 10% FBS in the presence of 5% CO_2_ at 37°C. Transfections used 500 ng of an HIV-1 infectious molecular clone (LAI, LAI SLQ^M^, T/F CH058, T/F CH470, T/F CH040, T/F CH077, T/F Thro, T/F CH236, T/F CH264, T/F CH850, or T/F CH569) and 25 or 50 ng of pVIVO2 containing No A3, A3G-HA, A3F-V5, or both A3F-V5 and A3G-HA. The pVIVO2 (Invivogen) has two MCS to enable co-expression of two genes on a single-cell basis. GeneJuice (EMD Millipore) transfection reagent was used according to manufacturer’s instructions. At 16 h post transfection the media was changed. Culture supernatants containing the virus were harvested 46 h post transfection, filtered through 0.45 μm polyvinylidene difluoride (PVDF) syringe filters and used to infect TZM-bl cells. TZM-bl cells were plated at 1 x 10^4^ cells per well of a 96-well plate and infected with a serial dilution of the virus in the presence of polybrene (8 μg/mL). The TZM-bl cells were maintained in DMEM with 10% FBS in the presence of 5% CO2 at 37°C. Forty-eight hours after infection the cells were washed with PBS and infectivity was measured through colorimetric detection using a β-galactosidase assay reagent and spectrophotometer. Infectivity of each virus was compared by setting the infectivity of the “No A3” condition as 100%.

### Immunoblotting cell and virus lysates

The cells and culture supernatants used for immunoblotting were obtained from the single-cycle infectivity assays. Cells were washed with PBS and lysed with 2x Laemmli buffer. Total protein in the cell lysate (30 µg) was resolved by SDS-PAGE for immunoblotting on a nitrocellulose membrane. A3 encapsidation was determined by concentrating the virus containing supernatant using Retro-X Concentrator (Clontech) following the manufacturer’s protocol, resuspending the pellet in 45 µL of Laemmli buffer and resolving 12 μL of the virus concentrate by SDS-PAGE for immunoblotting on a nitrocellulose membrane. HA tagged proteins were detected in cell lysates using mouse monoclonal anti-HA antibody (Cat# H9658, Sigma, 1:10,000). HA tagged proteins were detected in virus lysates using rabbit polyclonal anti-HA antibody (Cat# H6908, Sigma, 1:1,000). The V5 tagged proteins were detected in cell and virus lysates using anti-V5 mouse monoclonal antibody (Cat# V8012, Sigma, 1:1,000). Loading control for cell lysate was detected with anti-α-tubulin (rabbit polyclonal, Cat# PA1-20988, Invitrogen, 1:5,000) and for virus lysate was detected with anti-p24 (mouse monoclonal, Cat #3537, NIH HIV Reagent Program, 1:1,000). Secondary detection was performed using Licor IRDye antibodies produced in goat (IRDye 680-labeled anti-rabbit 1: 10,000 Cat # 926-68071 and IRDye 800-labeled anti mouse 1:10,000 Cat # 926-32210).

## Acknowledgments

We acknowledge Quinlan Carter for technical assistance with this work.

## Supporting information

**S1 Fig**. **HIV-1 T/F viruses have variable infectivity and Vif function in the presence of A3F, A3G, and A3F/A3G.** 293T cells were co-transfected with No A3 (empty vector), A3G-HA (A3G), A3F-V5 (A3F) or A3F-V5/A3G-HA (A3F/A3G) and an HIV-1 infectious molecular clone (labeled on figure). **(A)** The infectivity was determined relative to No A3 being set at 100% (hatched line). Error bars represent the error range from two independent experiments. Data was replotted from Fig 5 to compare to other HIV-1 T/F viruses. **(B-G)** For immunoblots, cell lysates and virus lysates were collected at 48 h resolved by SDS-PAGE and transferred to nitrocellulose membrane for probing with anti-V5 and anti-HA. The anti-α-tubulin and anti-p24 served as loading control for cell and virus lysate, respectively. The relative infectivity was determined using relative β-galactosidase activity from infected TZM-bl cells normalized to the No A3 condition.

