## Supplementary material for "Variability in HIV-1 Transmitted/Founder Virus Susceptibility to Combined APOBEC3F and APOBEC3G Host Restriction": S1 Fig

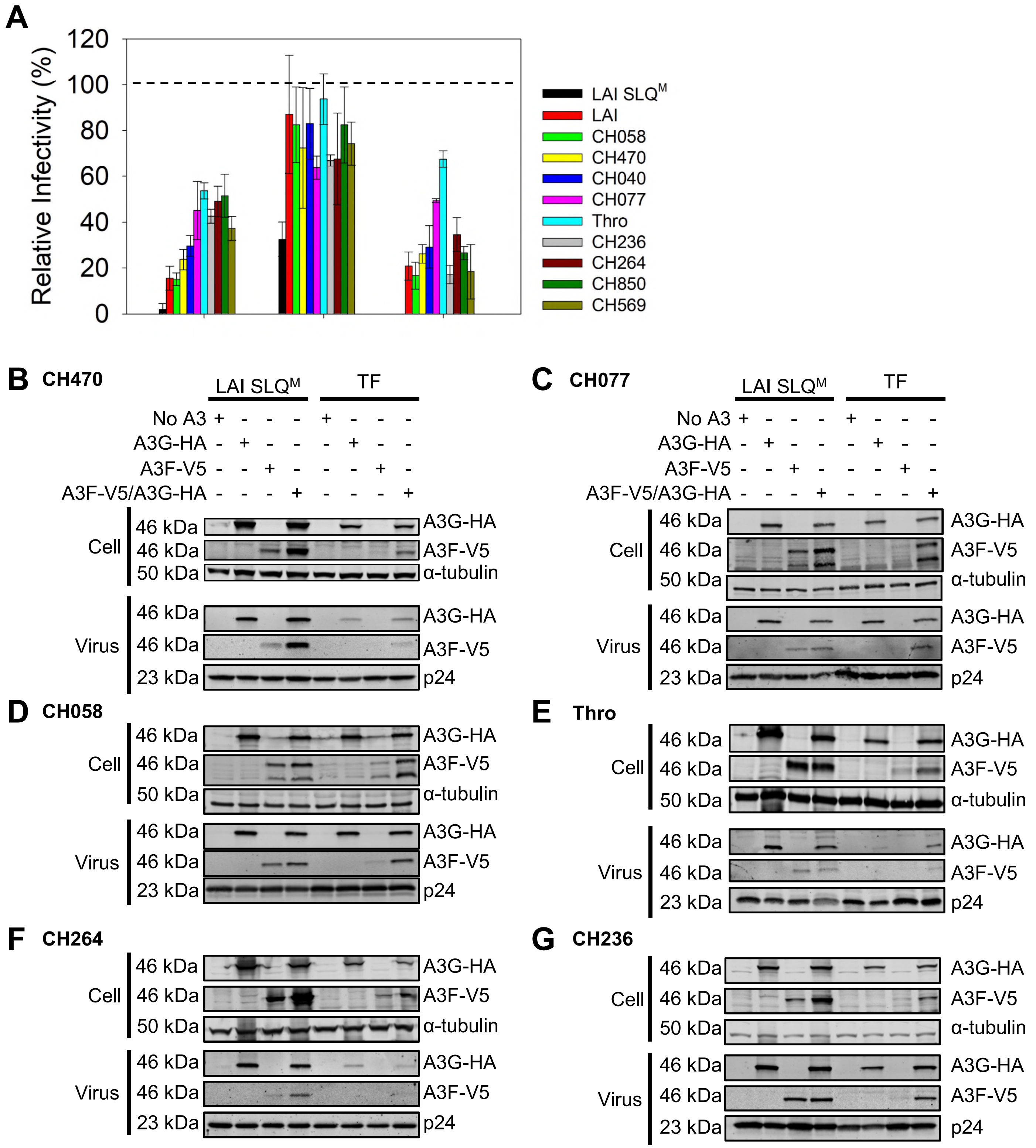
A
Relative Infectivity (%)
120
100
80
60
40
20
0
LAI SLQ^M
LAI
CH058
CH470
CH040
CH077
Thro
CH236
CH264
CH850
CH569
B CH470
C CH077
LAI SLQ^M
TF
No A3
A3G-HA
A3F-V5
A3F-V5/A3G-HA
Cell
Virus
46 kDa
50 kDa
23 kDa
α-tubulin
p24
D CH058
E Thro
F CH264
G CH236
